# Assessing the impact of mindfulness attitudes on attentional quality through focused breath induction

**DOI:** 10.1101/619817

**Authors:** M França, GK Tanaka, M Cagy, P Ribeiro, B Velasques, OL Gamboa

## Abstract

*Mindfulness attitudes*, as gentleness, openness, acceptance, curiosity and being non-judgmental have been related to improvement in cognitive and emotional functions, but few studies have focused on its specific contribution. The present study investigated the effect of the *mindfulness attitudes* on top-down attentional control abilities. Twenty one healthy participants were submitted to two conditions: a Mindfulness induction session where participants practiced focusing on the sensory sensations of breathing while encouraged to incorporate the five *mindfulness attitudes* and an attentional control session in which participants were repeatedly instructed to merely attend to the breath, without any mindfulness attitude component. Before and after each condition, participants performed two blocks of the oddball task while EEG was recorded. Contrary to our expectations, attentional control assessed through amplitude and latency of the P3b ERP component and oddball task accuracy did not show any changes in any of the conditions. These results suggest that a low dose of mindfulness training in naive individuals, through a focused breath induction, is not enough to improve the allocation of attentional resources towards task-relevant stimuli.

## 1. Introduction

Mindfulness has been described as a process of bringing a certain quality of attention to moment-to-moment experience (Kabat-Zinn, 1990). Mindfulness-based intervention (MBI) has been widely used to improve psychological morbidity associated with chronic illnesses and treat emotional and behavioral disorders (Cullen, 2011; Shonin, Van Gordon, & Griffiths, 2013).

In an effort to establish the defining criteria of its various components and specify implicated psychological processes, Bishop et al. (2004) have defined mindfulness as an attentional process taking place during the immediate experience, that involves the self-regulation of attention that leads to non-elaborative awareness and a display of traits such as curiosity, experiential openness, and acceptance. Further, the practice entails the development of a non-judgmental attitude toward one’s own experiences (Kabat-Zinn, 2006; Malinowski, 2008). Based on this operational definition we can argue that what makes mindfulness a particular attentional strategy is the interaction between the object of attention and the above mentioned set of *mindfulness attitudes* namely, curiosity, experiential openness, acceptance, and non-judgment (Dunn, Hartigan, & Mikulas, 1999; Ju & Lien, 2016; Teper & Inzlicht, 2014).

Despite the growing popularity of MBI, and the efforts to develop an operational definition of mindfulness (Bishop et al., 2004), there is still a general lack of agreement about what encompasses this quality of attention and the specific effect and relevance of the mindfulness components (Chiesa, 2013). The elusiveness of the concept of mindfulness as well as the varied ways in which it has been operationalized in research (Hart, Ivtzan, & Hart, 2013; Khoury et al., 2017) have made challenging to disentangle the psychological characteristics specific to mindfulness training from those shared with other experiential training approaches, such as, transcendental or concentrative meditation, metacognitive attentional training, or relaxation training (Bing-Canar, Pizzuto, & Compton, 2016; Nassif & Wells, 2014).

Few studies have focused on investigating the specific contribution of the *mindfulness attitudes* to the improvement in cognitive and emotional functions related to mindfulness, (Jo, Schmidt, Inacker, Markowiak, & Hinterberger, 2016; Lin, Fisher, Roberts, & Moser, 2016). There is evidence showing that being mindfully attentive with an attitude of curiosity operate in tandem to reduce defensive responding to a worldview threat. Mindful people show a willingness to consider new information about themselves and their world without reflexive judgements. In the absence of curiosity, mindfully attentive people appeared to be defensive, rejecting ideas and disparaging people that challenged the notion of human uniqueness (Kashdan, Afram, Brown, Birnbeck, & Drvoshanov, 2011). These findings indicate the importance of understanding better the effect of the elements that entail mindfulness.

One possible reason for the lack of research in this direction is that most of the experimental manipulation that tried to elucidate the acute neurophysiological mechanism of mindfulness has compared this quality of attention with “opposed” strategies as rumination, worry, unfocused attention, or psychoeducation (Arch & Craske, 2006; Broderick, 2005; Eddy, Brunyé, Tower-Richardi, Mahoney, & Taylor, 2015; Lai, MacNeil, & Frewen, 2015). Although valuable insight into the mechanism of mindfulness has been provided by these studies, their choice of control conditions has limited the possibility to understand the contribution and specific role of each of the *mindfulness attitudes* (Rahl, Lindsay, Pacilio, Brown, & David Creswell, 2017). For example, one can not be sure if a significant part of the effect reported in these studies resulted solely from the strengthening of attentional effort by mindfulness induction or if it has a contribution of any other component (Fan & Posner, 2004; Sarter, Gehring, & Kozak, 2006).

Recent research began exploring the role of *mindfulness attitudes* in more depth, using Event-Related Potentials (ERP), a neurophysiological measure of voltages generated in the brain structures in response to specific events or stimuli (Luck, Woodman, & Vogel, 2000). Dispositional mindfulness has been related to electrophysiological correlates (ERP components) of more efficient selective attention and discrimination (N100), as well as greater conflict monitoring (N200) and inhibitory control (P300) (Quaglia et al., 2016). Importantly, this relation remained significant after controlling for attentional control (through the attentional control scale), a possible confounding factor of mindfulness operationalization (Moore et al., 2012).

Teper and Inzlicht (2013) reported that experience in meditation enhances acceptance of emotional states, supposedly promoting better error perception and error reaction that led to better executive control. These results were indexed by amplification of the Error-Related Negativity component (ERN), a neural signal of error processing.

The present study aims to complement previous findings, investigating the effect of the *mindfulness attitudes* in the attentional quality. To achieve this, an oddball paradigm in combination with a focused breath induction exercise – an analog of mindfulness, to assess the effects of a first-time instruction (Arch & Craske, 2006) – were used to evaluate brain changes in the P3b wave. This ERP component can provide a measure of top-down attentional control abilities including the endogenous allocation of attentional resources towards task-relevant stimuli (Polich, 2009).

As a novelty, our control condition was an attentional exercise on the same object (breath) but without including any of the *mindfulness attitudes*. We reasoned that choosing the same object of attention could provide more straightforward information about the influence of the *mindfulness attitudes* on attentional quality. As mindfulness is recognized to improve the control of task-unrelated thought (Barron, Riby, Greer, & Smallwood, 2011) and strengthen the engagement of attention in the relevant stimulus (Atchley et al., 2016), we tested whether this effect would be higher when compared with a simple induction of attentional effort. As blinding participants in mindfulness studies have been a fundamental methodological problem (Davidson & Kaszniak, 2015), the similarity between our control and intervention conditions applied to a sample of naïve participants made possible a triple blind procedure.

We hypothesized that if the participants could minimally embody the *mindfulness attitudes* during the focused breathing induction procedure, it would cause attention improvement, compared with the attentional condition without *mindfulness attitudes.* This improvement would be expressed by a high amplitude of the P3b, as well as, better accuracy in the task.

## 2. Materials and methods

### 2.1 Participants

Twenty one healthy Participants (11 males, 28–39 years old, *M* = 37, *SD* = 3.8) were recruited from the Rio de Janeiro metropolitan area utilizing internet-based and word-of-mouth strategies, to be submitted to two conditions (mindfulness induction and control). All Participants were healthy, highly educated, right-handed (evaluated by the Edinburgh handedness inventory. Veale, 2014) and had normal or corrected to normal vision and naïve to meditation or correlated practices (e.g.Mindfulness, Yoga, Tai chi, martial arts, qi gong). Participants were not included if they had less than 6–8 h of sleep before the experiment and drunk caffeine during the 24 h prior to the experiment (Lorist & Tops, 2003).

The entire experimental protocol was explained to all participants who gave their signed consent before participating in this study. Five participants were excluded from all analyses: one participant fell asleep during the experiment and four participants presented electrode failure and/or excessive artifact at centro-parietal site. Thus a total of sixteen participants were included in the analyses reported in this study. This experimental protocol was approved by the Ethics Committee at the Federal University of Rio de Janeiro, Brazil

### 2.2 Procedure

Order of study procedures was identical between the focused breathing induction and control condition as showed in Figure 1. Participants initially completed a set of mindfulness questionnaires (e.g., MAAS, FFMQ) followed by placement of the EEG recording net. The experiment was performed in a sound and light-attenuated room, to minimize sensory interference. Participants were seated in a comfortable chair to reduce muscular artifacts while electroencephalography (EEG) data were collected. During the visual task, lights were turned off, and subjects were instructed to concentrate exclusively on the monitor screen. A 15” Samsung monitor was placed 50 cm in front of the participant, and they were submitted to one block of oddball task. After the task, participants completed a focused breathing induction exercise or control exercise followed by one more block of the oddball task. We adopted this pre-post protocol primarily to reduce the levels of the between-subjects factors. Such a design is particularly important for ERP studies, given the high degree of variability in the scalp-recorded EEG signals between individuals (Teper & Inzlicht, 2014). At the end of the experiment, participants were scheduled for the next visit (at least one week later). Half of the participants were randomly assigned to the intervention or attentional control condition in the first visit, to avoid order effect.

**Figure 1.**
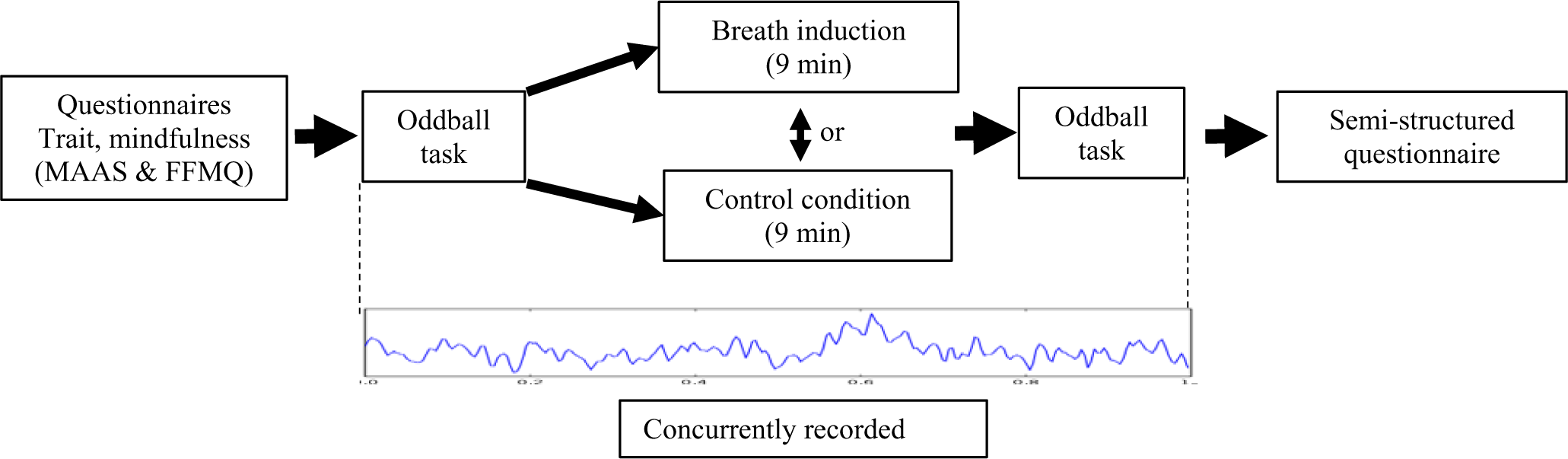
Overview of experimental procedure.

### 2.3 Mindfulness Individual Differences Measures

To account for potential differences in attention to the stimuli between participants based on the mindfulness-based trait, the Brazilian version (V. V. de Barros, Kozasa, Souza, & Ronzani, 2015; V. V. Barros, Kozasa, Souza, & Ronzani, 2014) of two trait mindfulness scales [Mindful Attention Awareness Scale (MAAS), and the Five Facet Mindfulness Questionnaire (FFMQ)] were administered at the first and second visit (Baer et al., 2008; Brown & Ryan, 2003).

#### Mindful Attention Awareness Scale

The MAAS is a 15-item questionnaire assessing trait aspects of mindfulness. The items are rated on a scale from 1 (almost always) to 6 (almost never). The total score is computed by taking the average of the responses to the 15 items. Higher scores reflect higher levels of dispositional mindfulness.

#### Five Facet Mindfulness Questionnaire

The FFMQ is a 39-item questionnaire also assessing trait mindfulness on five subscales: Observe (eight items) higher scores = more observant (highest possible score = 40), Describe (eight items) higher scores = more descriptive (highest possible score = 40), Act with Awareness (eight items) higher scores = more aware of actions (highest possible score = 40), Non-judge (eight items) higher scores = less judgmental (highest possible score = 40) and Non-react (seven items) higher score = better able to not react (highest possible score = 35). Each item is rated on a scale from 1(never or rarely true) to 5(very often or always true). Each subscale score is computed by summing the ratings for each item.

### 2.4 Mindfulness focused breathing induction and control condition

The recorded instructions for the focused breathing induction were adapted from the sitting mindfulness meditation exercise used by Kabat-Zinn (Mindfulness-based stress reduction) and had been reproduced in other studies (Arch & Craske, 2006; Yusainy & Lawrence, 2015).

In the Mindfulness induction condition, participants listened to an 8 min 30-second recording instructing them to establish a straight upright sitting posture, hands resting on their lap, shoulders relaxed, head upright, and feet resting flat on the floor. They were asked to keep their eyes open but without focusing on an external object, in particular, directing their attention to their internal experience. Participants were guided through instructions and engaged in the practice of focusing on the sensory sensations of inhalation and exhalation. They were also instructed to be fully present in the moment, to bring their attention back to the sensation of breathing when their mind wandered while cultivating the five *mindfulness attitudes:* gentleness, openness, acceptance, curiosity and being non-judgmental.

For the control condition, participants were repeatedly instructed to merely attend to the breath, without any mindfulness attitude component. Both recordings used the same voice and were balanced keeping the same time and amount of instruction to avoid facilitation by external feedback in favor of any condition. After all the procedures, participants were asked if they understood and were able to follow the instructions of the recording and if they stayed awake during the different parts of the session.

### 2.5 Oddball Task

All subjects were presented with the same visual discrimination task, which used the classical visual oddball task containing non-Kanizsa and Kanizsa figures (Illusory contours induced by “pacmen” image) as stimuli (Flynn et al. 2009). In this paradigm, four stimuli: two are kanizsa (internal triangle and square) and two non-Kanizsa (external triangle and square) figures are presented randomly, one of which occurs infrequently. Participants were asked to discriminate targets (25% infrequent) from non-targets or standard stimuli (75% frequent). Target stimuli were defined as internal squares. Participants were instructed to respond as quickly as possible only to target stimuli by pressing a button with their right index finger using a joystick (Quick Shot-Crystal CS4281, Quick shot, USA). Each participant received one block of 400 trials before and after the intervention, where specifically 100 target stimuli were presented in the block. Each stimulus appeared on the screen for 250 milliseconds, with inter-trial intervals (ITI) varying in the range of 1100–1300 ms. A fixation point (cross) was apresented during ITI (Figure 2).

**Figure 2.**
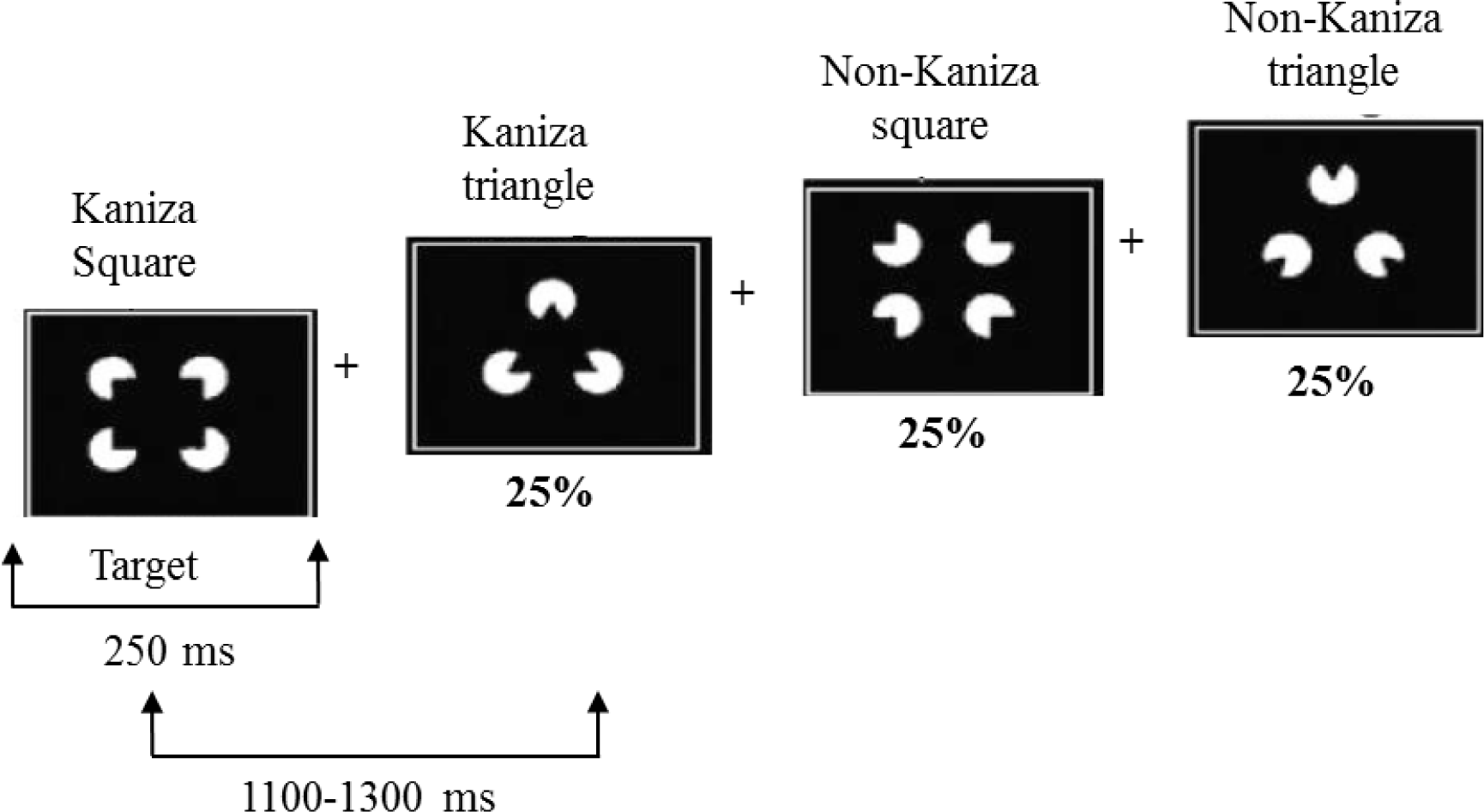
Kanizsa and non-Kanizsa figures as stimulus material. In_particular, the stimulus types are Kanizsa square (target), Kanizsa triangle, non-Kanizsa square, and non-Kanizsa triangle.

### 2.6 Electroencephalography data acquisition

EEG data were collected using a 20-channel nylon cap from the following locations: Fp1, Fp2, Fpz, F3, F4, F7, F8, Fz, C3, C4, T7, T8, Cz, P3, P4, P7, P8, Pz, O1, and O2 (Braintech-3000, EMSA-Medical Instruments, Brazil. Electro Cap Inc., Fairfax, VA, USA), yielding monopole derivations referred to linked earlobes. The EEG signal was amplified with a gain of 22,000, analogically filtered between 0.01 Hz (high-pass) and 100 Hz (low-pass). The software ERP Acquisition (Delphi 5.0), developed at the Brain Mapping and Sensorimotor Integration Laboratory, was employed to deliver the visual stimulus and record electrophysiological signals.

### 2.7 Blinding and randomization procedures

To ensure experimenter blinding to condition, an independent research-staff member created a pre-randomized set of labeled audio files for each participant and delivered record instructions by headphones. Subsequently the collected data was divided in two groups (control and mindfulness) and different codes were assigned to each group, keeping the research blinded during data processing and analysis.

The effectiveness of participants’ blinding was confirmed verbally at the end of each condition by asking if they knew what mindfulness is and if they associated any recording with it.

### 2.8 Behavioral data analyses

Differences in accuracy and reaction times were assessed using a 2×2 repeated measures ANOVA (rmANOVA), with factors Condition (mindful/control) and Time (pre/post). Normality of the data was verified using Shapiro-Wilk test. Sphericity was assumed in agreement with Mauchly’s sphericity test. Greenhouse-Geisser correction was used when sphericity was violated. The significance level was set at *p* ≤ .05. Statistical tests were performed using SPSS version 23.0 (SPSS Inc, Chicago).

### 2.9 EEG data processing and analysis

EEG data were preprocessed and analyzed offline using MATLAB 5.3 (Mathworks, Inc.) and EEGLAB toolbox (http://sccn.ucsd.edu/eeglab). To reduce line noise contamination a 60 Hz notch filter was applied. Data were first downsampled to 200 Hz and visually inspected to get rid of contaminated data due to nonneuronal activity such as muscle activity, electrode malfunction, or abrupt head movements. Data were filtered using a 0.1Hz high pass filter, 30 Hz low pass filter and impedances were kept below 10 kΩ. Independent Component Analysis (ICA) was then applied to identify and remove independent components resembling eye-blink or muscle artifact. Artifact-free EEG data of the correctly responded and non responded trials, were re-referenced to an average reference and segmented from −200ms before to 1000 ms after stimulus presentations. Segments were baseline corrected using a 200 ms pre-stimulus epoch and averaged. Since our primary measures were the attentional P3b ERP component, specifically in response to infrequent target, latency and amplitude were quantified in the time window from 320 to 600 ms, analyzed at the Pz electrode site. No between – Condition differences were observed regarding the number of trials available for analysis (Mindfulness condition: *M* = 76.6, 78.8; *SD* = 12.5, 12.3; Control condition: *M* = 76.6, 77.2, *SD* = 11.8, 13.8, for pre and post interventions, respectively) or the amount of independent components (Mindfulness condition: *M* = 18.6, 18.4, *SD* = 0.5, 0.6; Control condition: *M* = 18.9, 18.8, *SD* = 0.2, 0.3, for pre and post interventions, respectively).

The relationship between the experimental manipulations and brain activity was examined for the preselected P3b component at Pz, through a 2 × 2 repeated measures ANOVA (rmANOVA), with factors Condition (mindful/control) and Time (pre/post). Sphericity was tested with Mauchly’s test of sphericity, and Greenhouse-Geisser correction was applied if necessary. The significant threshold was set at *p* ≤ .05. Statistical tests were performed using SPSS version 23.0. (SPSS Inc, Chicago).

### 2.10 Relating Trait Mindfulness Individual Differences to ERP Effects

Because we were interested in how individual differences in trait mindfulness related to ERP effects, we correlated mean amplitude and latency P3b with the MAAS and FFMQ overall scores. With exploratory purpose, a correlational analysis between the p3b component and FFMQ subscale was conducted.

## 3. Results

### 3.1 Behavioral results

Evaluation of the performance in the Oddball task showed no difference in reaction time, in the factor Condition *F* (1,15) = 1.12, *p* = 305 and Time *F* (1,15) = .328, *p* = 578. Analisys of commission error to Condition *F* (1,15) = 3.87, *p* = .068 and Time *F* (1,15) = .242, *p* = .630, did not reach significance (breath induction pre 421 ms and post 422 ms, control condition pre 419 ms and post 422 ms. See Figure 3a). Response accuracy (percent of correct hits) in the factors Condition *F* (1,15) = .044, *p* = .837 and Time *F* (1,15) = 1.06, *p* = .319, was not significant (breath induction pre 99,7% and post 99,6%, control condition pre 99,6% and post 99,4%. See Figure 3b). We also evaluated miss response, that revealed no statistical significance to Condition *F* (1,15) = 0.44, *p* = 837 and Ttime *F* (1,15) = 1.06, *p* = .319. There were no intertaction between Condition and time.

**Figure 3.**
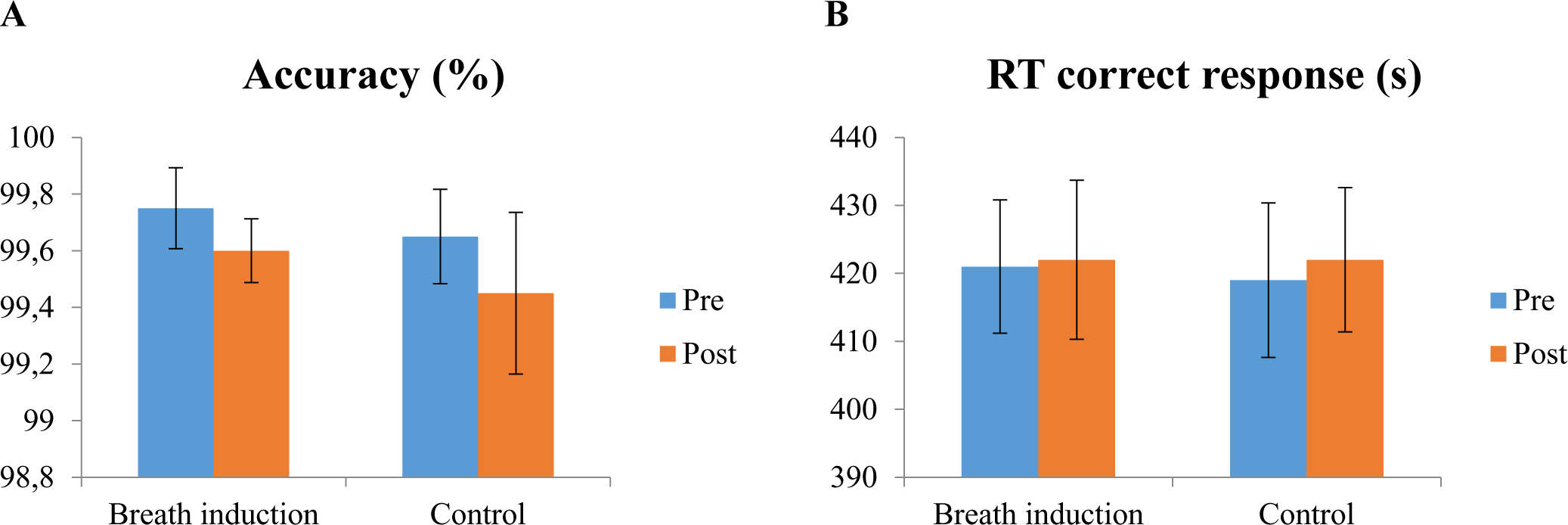
Mean reaction time (RT) and accuracy of the Oddball task. Pre and post breath induction and control condition. Errors bar represent standard errors.

### 3.2 ERP Results

Electrophysiological data, including the mean amplitude and latency of the factors and Time, are presented in Table1. The grand average of the ERPs at the pz site and topographic maps are shown in Figure 4.

**Figure 4.**
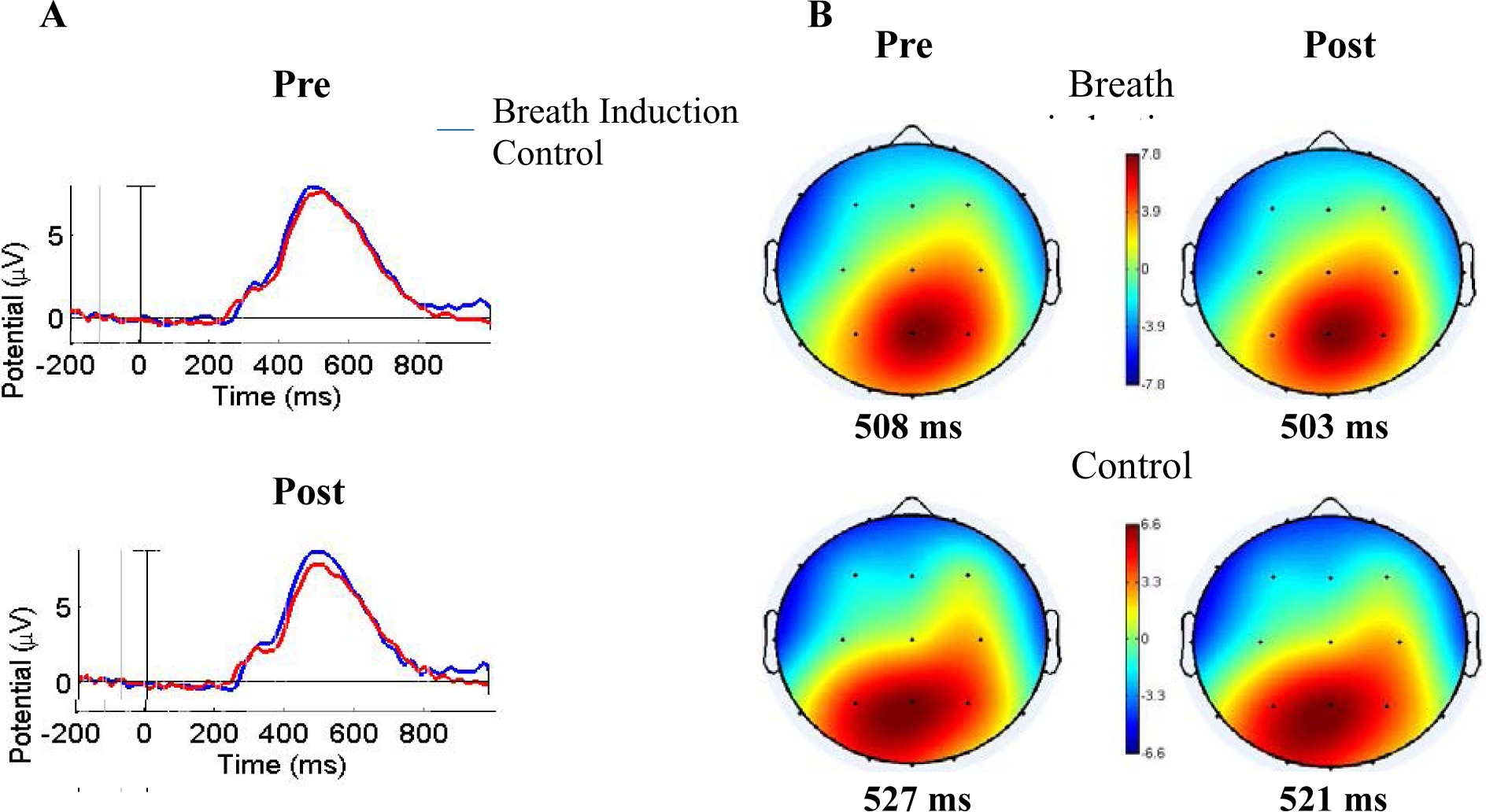
Grand averages ERPs and voltage maps for the P3b component at the electrode Pz for both conditions (n=16). **A.** Grand average P3b showing the average in the centro-parietal electrode (Pz) for control and breath induction conditions, pre (left) and post (right). **B.** Pre and post voltage maps at the peak latency of the P3b component for both conditions. Maps are individually scaled and color-coded in agreement with their respective maximum (red) and minimum (blue).

**Table 1.** Mean and Standard Deviation (SD) of the P3b ERP component.

| P3b<br>320–600 ms | Breath induction condition |  |  |  | Control condition |  |  |  |
| --- | --- | --- | --- | --- | --- | --- | --- | --- |
|  | pre |  | post |  | pre |  | post |  |
|  | mean | SD | mean | SD | mean | SD | mean | SD |
| Amplitude (μV) | 8.0 | 2.22 | 8.31 | 2.56 | 7.94 | 3.16 | 8.20 | 2.56 |
| Latency (ms) | 508 | 40.31 | 503 | 31.17 | 527 | 48.68 | 521 | 45.70 |

To assess the effect of the mindfulness focused breath induction, the Two-Way ANOVA compared condition (brief mindfulness induction, control) and time (pre and post-intervention) within the centro-parietal electrode.

Target: The factor Condition did not show significant differences (see Figure 4) in terms of amplitude (*F* (1,15) = .016, *p* = .901) or latencies (*F* = (1,15) = 1.70, p = .212). Similarly, amplitudes (*F* (1,15) = .688) and latencies (*F* (1,15) = 1.67, *p* = .216) were not significantly different in the factor Time (see Figure 4). There was no interaction between factors, when assessing amplitude *F* (1,15) = .007, *p* = .934 and latency *F* (1,15) = .012, *p* = .914.

Non Target: The factor Condition did not show significant differences in terms of amplitude (*F* (1, 15) = .069, *p* = .796) or latencies (*F* (1, 15) = 2.28, *p* = .152). Similarly, for the factor Time, amplitudes (*F* (1, 15) = .439, *p* = .518) and latencies (*F* (1, 15) = 0.93, *p* = .764) were not significantly different. Similarly, no interaction between factors was observed (amplitude *F* (1,15) = 2.99, *p* = .104 and latency *F* (1,15) = 3.81, *p* = .070).

### 3.3 Individual Differences in mindfulness trait and ERP Effects

We expected that individual differences in mindfulness trait would influence the pattern of ERP effects during the oddball task. Therefore, we examined the relationship between trait mindfulness (MAAS and FFMQ) and P3b amplitude and latency, pre and post induction and control. Our results did not show any significant correlation between P3b ERP component and Trait Mindfulness scales.

## 4. Discussion

The current study examined the impact of the *mindfulness attitudes*, through a brief mindfulness induction relative to ordinary attention, on behavioral and electrophysiological indices of cognitive control. Specifically, we sought to disclose if the attitudes of nonjudgement, acceptance, curiosity, and openness could affect attentional control evidenced by modulation of the p3b component and task accuracy. Contrary to our hypotheses, our results showed that a brief mindfulness induction did not cause an alteration in P3b component and task performance when compared to the control task. These results reflect that one session of a focused breath induction exercise in naïve participants is not enough for them to learn how to embody the *mindfulness attitudes*. Besides, due to the characteristic of the control task, it is feasible that participants in both conditions used similar attentional strategies, leaving no room for significant differences.

We are not aware of any previous study that used only one dose of mindfulness induction procedure and the P3b as an electrophysiological marker of attentional quality. However, other ERP components related to cognitive control have been investigated. Resembling our results, studies that used only one dose of training in naïve subjects did not find any influence on performance-related indices of cognitive control. For example, Larson et al. (2013), did not find significant differences between groups for behavioral performance or ERN amplitudes or latencies, after a mindfulness induction procedure following a 14-min audio clip focused on attending to their breathing and being mindful of the moment (Larson, Steffen, & Primosch, 2013). Alike Bing-Canar et al., (2016) using the same mindfulness induction failed to reach significance for the ERN but found changes in the alpha power activity and enhanced error-related alpha suppression during the subsequent Stroop task. These modulations reflect a mental state that is relaxed or characterized by the inward focus of attention and is correlated to greater attentional engagement following errors compared to correct responses (Bing-Canar et al., 2016). Conversely, when naive participants underwent an induction procedure that emphasized being mindful of present-moment feelings during an emotional go/no-go task, ERN magnitude was increased. The authors argue that when the brief mindfulness inductions is specially adapted, can be powerful enough to influence ERN amplitudes (Teper & Inzlicht, 2014). All these different electrophysiological modulations may suggest that some measures are more sensitive (e.g., alpha power and occasionally ERN) or related to cognitive aspects that change very early during mindfulness training.

Previous attention research performed with experienced meditators has shown the sensitivity of the P3b component to several levels of mindfulness training (Kaunhoven & Dorjee, 2017, Delgado-Pastor, Perakakis, Subramanya, Telles, & Vila, 2013). Therefore, our lack of P3b modulation could indicate that the processes associated to theP3b component undergo changes during more advanced stages of the practice.

It has been reported that P3b amplitude is sensitive to task difficulty (Polich, 2009), mindfulness training may increase attention flexibility and adjust attention resource allocation depending on task demands (Kaunhoven & Dorjee, 2017).The task accuracy rate in our study was very high, with all participants showing almost maximum accuracy (mean of 99 correct responses of a total of 100 targets, pre-post in both conditions). It is possible that the easy nature (based on our participant’s skills) of our oddball task was insufficient to cause significant changes in the process underlying the P3b or uncover any modulation caused by *mindfulness attitudes*. This could also help to explain why we did not find a correlation between trait mindfulness (measured by MAAS and FFMQ), P3b, and task performance.

We are aware of only one study that investigated the relationship between dispositional mindfulness and attentional control during a discrimination face go/no-go task (Quaglia, Goodman, & Brown, 2016). The authors found that high dispositional mindfulness was associated with changes in ERP related with great inhibitory control (more negative N100), conflict monitoring capacity (more negative N200) and also predicted faster response time.

The implications of the present findings should be considered in light of some limitations. Due to the characteristic of the control task, it is feasible that participants used similar attentional strategies in both conditions, leaving no room for significant differences. Additionally, the high accuracy observed during task performance might indicated a ceiling effect that could have contributed to the lack of differences between conditions (Jo et al., 2015).

In conclusion, the present study suggests that one dose of mindfulness induction when compared to a common attention condition in the same object (breath) is not enough to cause an improvement of the attentional quality. We could not find any changes produced by *mindfulness attitudes*. We speculate that the absence in the modulation of task accuracy and P3b may be due to the low dose of mindfulness training, and lack of adequacy of the task: low level of task difficulty and lack of sensitivity to the P3b to early changes produced by mindfulness training. Further research with a similar study design but adopting a more demanding task and great exposure to training, is needed to confirm and extend the present negative findings.

## 5. Competing financial interest

The authors declare no competing financial interests

## 6. Acknowledgements

We thank Ivonne Isabel Fuentes for her help during various stages of the project and all the participants that took part in this study.

